# Distinct Roles of Somatostatin and Parvalbumin Interneurons in Regulating Predictive Actions and Emotional Responses During Trace Eyeblink Conditioning

**DOI:** 10.1101/2025.03.23.644831

**Authors:** Jiaman Dai, Qian-Quan Sun

## Abstract

Learning involves evaluating multiple dimensions of information and generating appropriate actions, yet how the brain assigns value to this information remains unclear. In this study, we show that two types of interneurons (INs) in the primary somatosensory cortex—somatostatin-expressing (SST-INs) and parvalbumin-expressing (PV-INs) neurons—differentially contribute to information evaluation during trace eyeblink conditioning (TEC). An air puff (unconditioned stimulus, US) delivered after a whisker stimulus (conditioned stimulus, CS) elicited both reflexive eye closure and stress-related locomotion. However, only self-initiated, anticipatory eye closure during the CS window, measured via electromyography (EMG), was directly relevant to learning performance. We found that SST-IN activity changes aligned with the learning induced changes of the anticipatory eye blinks during the CS period, correlated with the EMG changes across learning. In contrast, PV-IN activity was positively correlated with stress-related locomotion following the US and showed no learning related changes, suggesting a role in processing the emotional or aversive component of the task. Furthermore, cholinergic signaling via nicotinic receptors modulated both SST- and PV-IN activities, in a manner consistent with their distinctive roles, linking these interneurons to the regulation of learning-related actions and emotional responses, respectively. These findings demonstrate that distinct interneuron populations evaluate different dimensions of information—SST-INs for predictive, adaptive actions and PV-INs for stress-related emotional responses—to guide learning and behavior.

## Introduction

In the early stages of learning, the brain is bombarded with a diverse array of information— sensory inputs like touch or sound, contextual cues such as location, and emotional signals like fear or reward. Evaluating the relevance of this flood of data and generating appropriate responses are essential for adaptive behavior. Traditionally, the primary sensory cortex, the brain’s first stop for processing raw sensory information, was thought to handle only basic features, leaving higher-order cortical areas—regions responsible for complex integration—to guide decision-making and behavior(Felleman and Van Essen, 1991; Freedman and Assad, 2016). However, recent discoveries challenge this view, revealing that even the superficial layers of the sensory cortex encode contextual details during learning, such as self-motion, expectation, and attention (Ayaz *et al*., 2019; Makino *et al*., 2016; Poort *et al*., 2015). This expanded role suggests the sensory cortex is not just a passive relay but an active hub for integrating multimodal information critical to learning.

A defining feature of this cortical processing is the diversity of interneurons (INs), specialized cells that fine-tune neural activity through inhibition. Subtypes like somatostatin-expressing (SST-INs) and parvalbumin-expressing (PV-INs) interneurons play distinct roles based on their structure and connections(Letzkus *et al*., 2011; Pfeffer *et al*., 2013; Tremblay *et al*., 2016). SST-INs target dendrites—the branched extensions of neurons—to shape sensory integration and plasticity, while PV-INs target cell bodies to control network synchrony and amplify signals (Garcia-Junco-Clemente *et al*., 2019; Karnani *et al*., 2016; Kepecs and Fishell, 2014). Another subtype, vasoactive intestinal peptide-expressing interneurons (VIP-INs), adds further complexity by disinhibiting circuit (Ferguson *et al*., 2023; Myers-Joseph *et al*., 2024). Despite these well-characterized roles, how these interneurons contribute to the multifaceted nature of learning—spanning sensory prediction, emotional processing, and behavioral adaptation— remains a puzzle.

Building on this diversity, recent studies have begun to uncover how IN subtypes shape learning across different tasks and brain regions. For instance, in a lever-press task, VIP-INs in the motor cortex activate strongly during early learning, likely facilitating plasticity by silencing inhibitory brakes, while SST-INs show weaker responses (Szadai *et al*., 2022). In associative fear learning, PV-INs in the auditory cortex gate sensory inputs to sharpen fear memory formation (Letzkus *et al*., 2011), whereas in visual discrimination tasks, learning boosts PV-IN selectivity while SST-INs drift from local network patterns (Khan *et al*., 2018). In the prefrontal cortex, SST-INs fire during reward approach in cue-based tasks, while PV-INs signal reward departure, hinting at roles in encoding temporal sequences (Kvitsiani *et al*., 2013). A 2025 study further showed that lateral inhibition from SST-INs more effectively changed the gain of neural and perceptual contrast sensitivity(Del Rosario *et al*., 2025). These findings reveal that IN subtypes are not static players but dynamically engaged depending on the learning context and information type.

Yet, the role of IN subtypes in the primary somatosensory cortex (S1) during trace eyeblink conditioning (TEC)—a classic model of associative learning—remains uncharted territory. TEC pairs a conditioned stimulus (CS), like a whisker deflection(Dai and Sun, 2024), with an unconditioned stimulus (US), such as an air puff to the eye, separated by a brief trace interval. This task demands the integration of sensory, temporal, and emotional information, making it an ideal lens for studying IN contributions(Moyer *et al*., 1990; Woodruff-Pak, 1993). While the hippocampus and prefrontal cortex bridge the trace interval (McEchron *et al*., 1998) and the cerebellum times the conditioned blink (Christian and Thompson, 2005), the auditory cortex’s role in similar paradigms suggests S1 might also be key (Letzkus *et al*., 2011). However, no studies have probed how S1 interneurons encode sensory and contextual signals during TEC, leaving a critical gap in our understanding.

To fill this void, we tracked animal behavior and used two-photon imaging to monitor SST-INs and PV-INs in layer 2/3 of S1 during TEC. Our results reveal a striking division of labor: SST-IN activity changes aligned with anticipatory eye blinks during the CS, showing a negative correlation with electromyography (EMG) responses during TEC learning, while PV-IN activity tracked stress-related locomotion after the US and remain unchanged during TEC, suggesting a role in processing emotional valence. Cholinergic signaling via nicotinic receptors further modulated both subtypes, tying them to learning-related actions and affective states. This unexpected split—SST-INs driving predictive behaviors and PV-INs encoding stress—redefines S1’s role in associative learning. These insights, bolstered by recent evidence of IN dysfunction in psychiatric disorders(Gogolla *et al*., 2009; Marín, 2012), illuminate how cortical interneurons orchestrate the integration of sensory and emotional information. By decoding SST-IN and PV-IN roles in TEC, this study offers fresh perspectives on adaptive behavior and potential therapeutic targets for neuropsychiatric conditions like schizophrenia, autism, and anxiety, where interneuron imbalances are implicated(Gogolla *et al*., 2009; Marín, 2012).

## Results

### Changes in Interneuron Population Activity during TEC Learning

In our previous work, we demonstrated that the S1 is essential for TEC learning(Dai and Sun, 2024). To investigate how INs respond during TEC learning, we trained awake, head-fixed mice on a TEC paradigm while allowing them to move freely on a running wheel(Dai and Sun, 2024). In our setup, whisker deflection served as the conditioned stimulus (CS; 60 Hz, 250 ms), followed by an air puff to the cornea as the unconditioned stimulus (US; 20 psi, 50 ms) after a 250 ms trace interval (Fig. 1a). To isolate learning-induced changes, we used three stimulation patterns within each session: CS paired with US (16/20 trials, CS-US), CS alone (2/20 trials, CS-only), and US alone (2/20 trials, US-only). During these sessions, we imaged calcium transients (via GCaMP6s) in SST-INs or PV-INs through a 3 mm cranial window using two-photon microscopy. The region of interest (ROI) was identified through intrinsic imaging (Fig. 1b, c). Mice were awake and free to walk on a 3D-printed running wheel during imaging.

**Figure 1.**
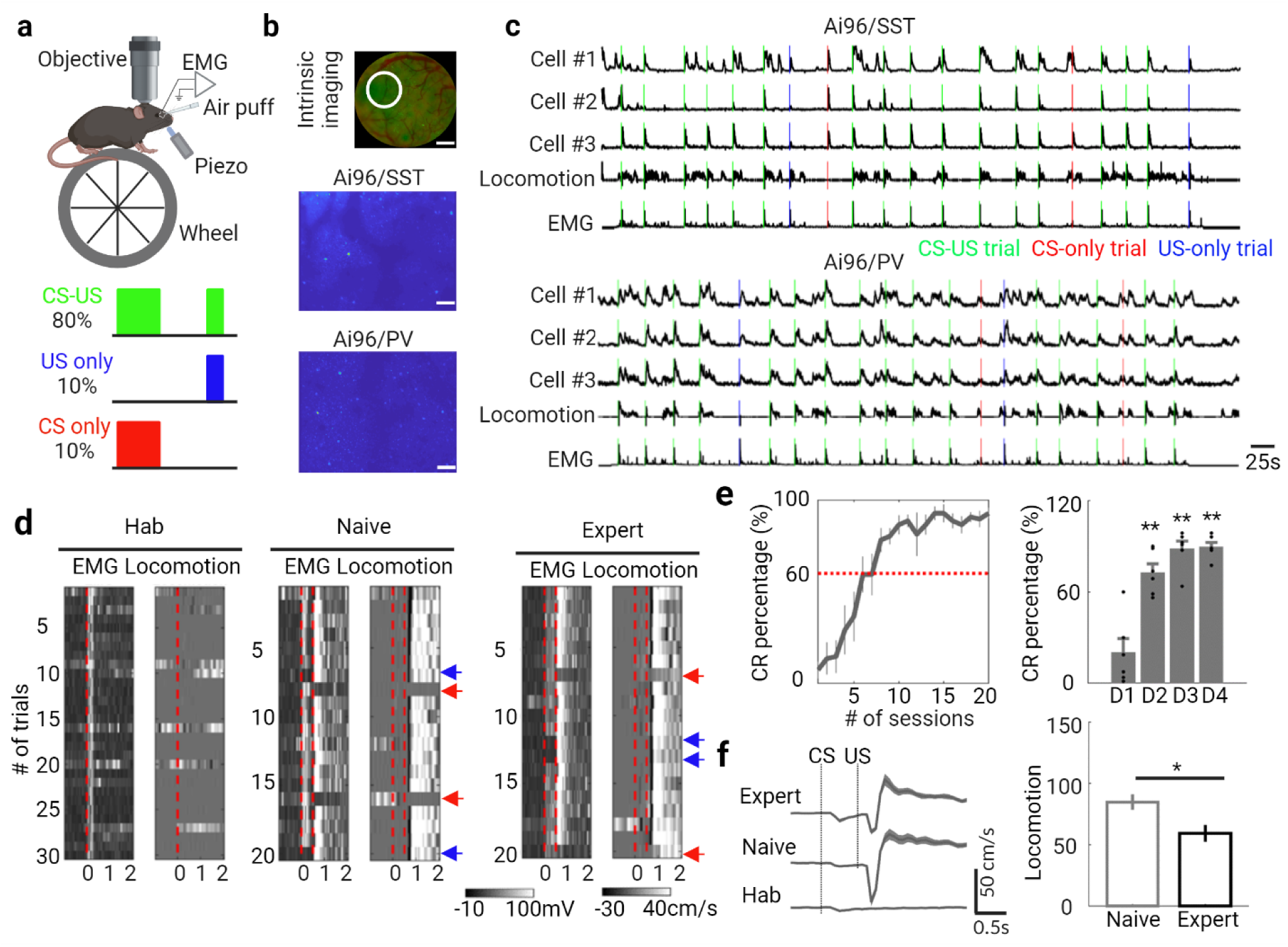
TEC learning task. **a.** Schematic representation of the experimental setup (top) and session structure (bottom), created with BioRender.com. **b.** Intrinsic imaging signals used to identify the whisker-evoked area in the barrel cortex (top, indicated by the white circle; scale bar, 500 µm). Example fields of calcium imaging for SST- and PV-INs in S1 are shown below (scale bar, 50 µm). **c.** GCaMP6s signal (Δf/f) traces from four example neurons, along with locomotion and EMG traces during the first session of TEC training (green: CS-US; red: CS-only; blue: US-only). **d.** EMG and locomotion activity during each trial at each training stage, with vertical dashed lines indicating the onset of CS or US. Orange arrows denote CS-only trials, blue arrows indicate US-only trials, while all other trials are CS-US. **e.** Evolution of the conditioned response percentage (CR %) across 20 sessions (top). Changes in CR percentage during training are shown below (n=12 mice; Day 1 vs. Day 2, **P=0.0087; Day 1 vs. Day 3, **P=0.0022; Day 1 vs. Day 4, **P=0.0022; Wilcoxon test). Data are expressed as mean ± s.e.m., with shaded areas representing s.e.m. **f.** Locomotion traces at different training stages, with the vertical dashed line indicating the onset of stimuli (first: whisker stimulation, CS; second: air puff, US). Notable changes in locomotion following US at both the naive and expert stages are observed (n=12 mice; *P=0.012; Wilcoxon test).

We focused on behavioral characteristics at three stages of TEC training we described in our earlier work(Dai and Sun, 2024): the habituation stage, where mice were exposed to unreinforced CS-only trials; the naive stage, representing the first reinforced session; and the expert stage, defined as the last session (Fig. 1d). After repeated CS-US pairings across multiple sessions, mice exhibited a gradual improvement in performance, reflected in increased orbicularis oculi muscle contractions (measured via EMG) during the interval between CS onset and US onset. The probability of conditioned eyeblink responses (CR %) increased progressively from 11% ± 5% to 86% ± 4% (Fig. 1e), consistent with previous studies from our group(Dai and Sun, 2024) and others (Giovannucci *et al*., 2017; Heiney *et al*., 2014). Additionally, locomotion following the US decreased from the naive stage to the expert stage (Fig. 1f).

Next, we examined IN activity in layer 2/3 (100–250 μm below the pial surface) of S1 across different training stages. We found that the average population activity of SST-INs was prominent during early CS and modulated by TEC training (Fig. 2a-c). Notably, GCaMP6 signals during the CS window (from CS onset to US onset, CR) increased from the habituation stage to the naive stage (0.36 ± 0.03 vs. 0.47 ± 0.04 ΔF/F, P = 6.16×10⁻⁹; Wilcoxon test) but decreased from the naive to the expert stage (0.47 ± 0.04 vs. 0.40 ± 0.04 ΔF/F, P = 1.43×10⁻⁶; Wilcoxon test). Similarly, calcium activity during the US window (1 s after US onset, UR) decreased from the naive to the expert stage (1.0 ± 0.08 vs. 0.68 ± 0.07 ΔF/F, P = 3.70×10⁻¹⁴; Wilcoxon test). Moreover, the correlation between calcium signals and locomotion decreased from the naive stage to the expert stage, while the correlation between calcium signals and EMG increased from the habituation to the naive stage (Fig. 2d). Thus, activities of SST-INs are correlated with anticipatory eye blinks during the CS period and correlated with the EMG changes across learning, and reduced after the animal already acquired the TEC. These results suggest that SST-INs behavior is an integral part of the S1 encoding of TEC learning.

**Figure 2.**
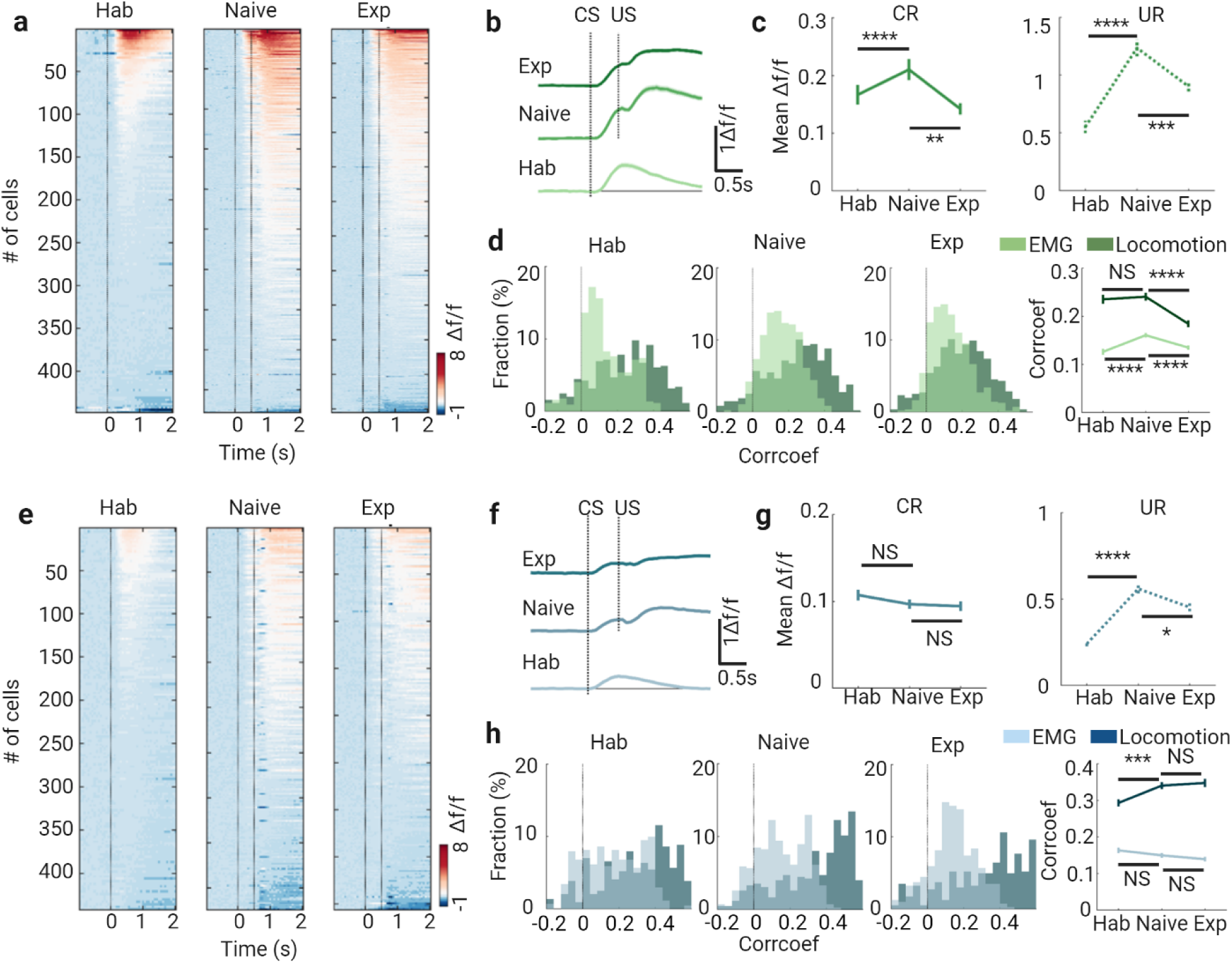
Changes in Population Activities of Interneurons during TEC Learning. **a.** Heat maps of GCaMP6s signals (Δf/f) for all somatostatin interneurons (SST) at different learning stages, aligned to the CS onset. **b.** Mean signal traces of SST at each learning stage, presented as mean ± s.e.m., with shaded areas representing s.e.m. **c.** Responses following CS delivery (conditioned response, CR; solid line, activity in CS window) and US delivery (unconditioned response, UR; dashed line, activity in US window) of SST at each learning stage (CR: ****P=1.12 × 10⁻⁷, **P=0.0064; UR: ****P=5.23 × 10⁻⁴⁶, ***P=4.10 × 10⁻⁴; Wilcoxon test). **d.** Distribution and changes in correlation coefficients between calcium signals and locomotion (dark green) / EMG (light green) at the three training stages (Locomotion: NS P=0.75, ****P=2.04 × 10⁻⁸; EMG: ****P=3.06 × 10⁻⁶, ****P=9.85 × 10⁻⁶; Wilcoxon test). **e.** Heat maps of GCaMP6s signals (Δf/f) for all parvalbumin interneurons (PV) at different learning stages, aligned to the CS onset. **f.** Mean signal traces of PV at each learning stage, shown as mean ± s.e.m., with shaded areas representing s.e.m. **g.** Responses after CS delivery (CR; solid line, activity in CS window) and US delivery (UR; dashed line, activity in US window) of PV at each learning stage (CR: NS P=0.94, NS P=0.99; UR: ****P=3.14 × 10⁻²², *P=0.011; Wilcoxon test). **h.** Distribution and changes in correlation coefficients between calcium signals and locomotion (dark green) / EMG (light green) at the three training stages (Locomotion: ***P=1.44 × 10⁻⁴, NS P=0.30; EMG: NS P=0.15, NS P=0.36; Wilcoxon test).

Interestingly, TEC modulated PV-INs differently. While no significant changes in CR were observed across the learning stages (Fig. 2g), the UR of PV-INs exhibited similar changes to those seen in SST-INs across the stages (Fig 2g). Additionally, no significant correlation changes between calcium signals and EMG were observed in PV-INs during TEC training (Fig 2h), which contrasts with the activities of SST-INs. The correlation between calcium signals and locomotion increased from the habituation to the naive stage, but no difference was noted between the naive and expert stages (Fig. 2h). This may indicate that SST-INs play a more prominent role in evaluating sensory stimuli during learning, while PV-INs exhibit more stable activity patterns.

### Differential Locomotion-Modulated Activity in SST- and PV-INs across Learning Stages

Locomotion is known to modulate sensory processing (Ayaz *et al*., 2019; Niell and Stryker, 2010). We examined how locomotion influences the activity of SST- and PV-INs at different stages of TEC training. Based on recorded running speed, we categorized trials into two behavioral states: running and resting. Trials with a maximum speed below 5 cm/s (measured within 250 ms before and after CS delivery) were classified as resting trials.

At the habituation stage, SST-IN activity was lower during running trials compared to resting trials, while PV-IN activity was higher in running trials than in resting ones (Fig 3a vs. e). At both the naive and expert stages, SST-IN activity showed no significant difference between running and resting trials, either in the CS or US windows. However, at the naive stage, PV-IN activity during running trials remained higher than during resting trials in both the CS and US windows (Fig 3b vs. f). As learning progressed, no significant differences in PV-IN activity were observed between running and resting trials in either window (Fig 3g). These findings suggest that SST- and PV-INs are differentially modulated by locomotion across learning stages, with PV-INs showing a greater sensitivity to movement during early learning. This may reflect distinct roles for these interneuron subtypes in integrating motor behavior with sensory processing during task acquisition.

**Figure 3.**
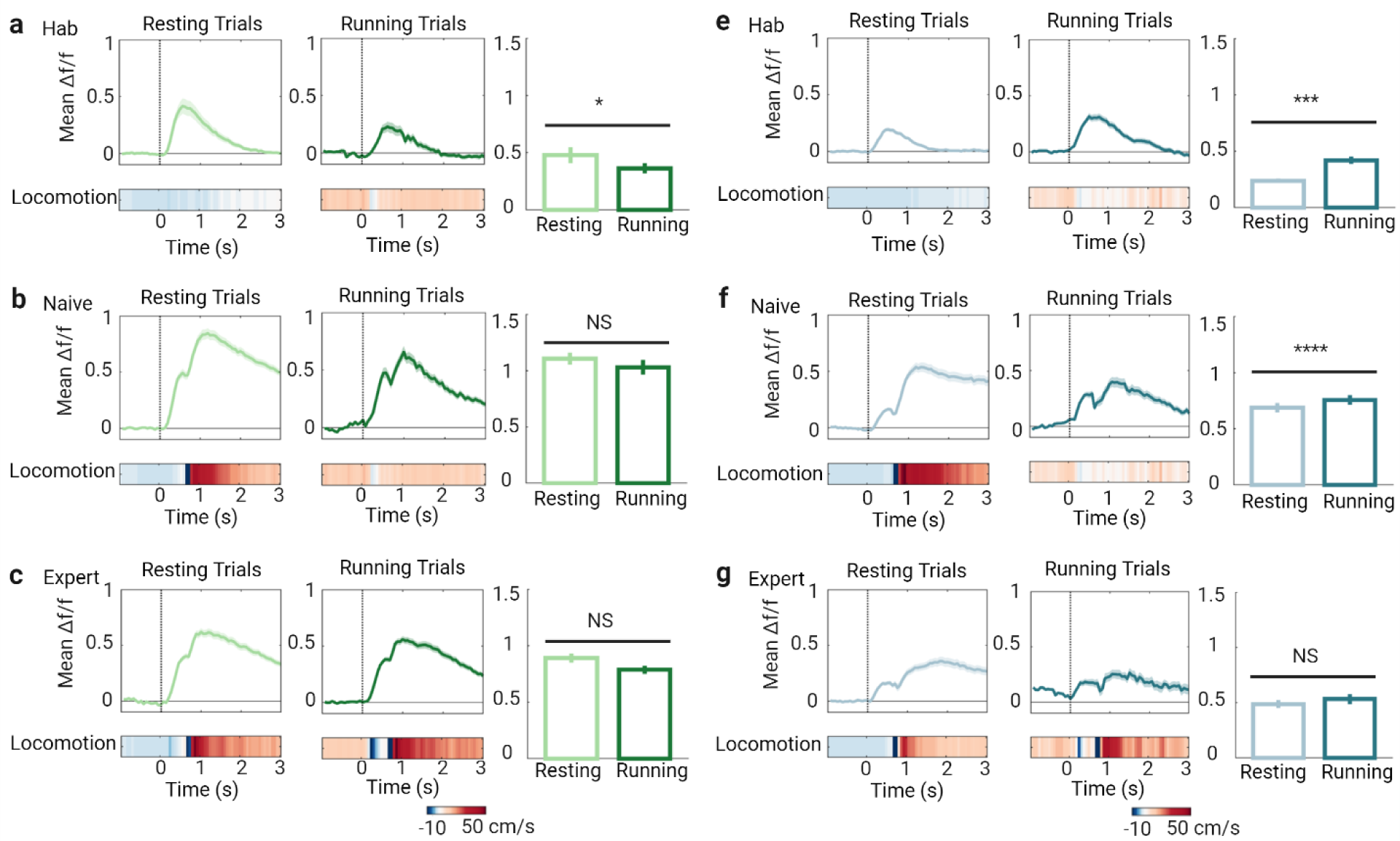
Locomotion-Dependent Activity in SST and PV Interneurons During TEC Learning. **a.** Mean signal traces of somatostatin interneurons (SST) during resting and running trials (left and middle top). The corresponding heatmap of mean locomotion is presented below (left and middle bottom), along with comparisons on the right at the habituation stage. Data are expressed as mean ± s.e.m., with shaded areas indicating s.e.m. **b.** Same as (a) but at the naive stage. **c.** Same as (a) but at the expert stage. **e.** Mean signal traces of parvalbumin interneurons (PV) during resting and running trials (left and middle top), with the corresponding heatmap of mean locomotion presented below (left and middle bottom) and comparisons on the right at the habituation stage. Data are shown as mean ± s.e.m., with shaded areas indicating s.e.m. **f.** Same as (e) but at the naive stage. **g.** Same as (e) but at the expert stage.

Combined with our previous observation that occurrence of locomotion were after UR period, and were much higher incidence during CS-US trials than CS-only trials (see also Fig 1c), and that locomotion was significantly reduced at the Exp. stage(Dai and Sun, 2024), these data suggest that locomotion is likely stress-related behaviors which are significantly reduced as animals again protection from anticipatory eye blinks. Thus, PV-INs activity was positively correlated with stress-related locomotion following the US and showed no learning related changes, suggesting a role in processing the emotional or aversive component of the task. These findings together suggest that SST-INs and PV-INs are differentially modulated during TEC learning, with SST-INs showing dynamic activity changes across learning stages. This may indicate that SST-INs play a more prominent role in evaluating sensory stimuli during learning, while PV-INs exhibit more stable activity patterns that are tuned to the emotional states.

### Cholinergic Modulation of TEC-Induced Neuronal Responses in Interneurons

Cholinergic modulations facilitate TEC conditioning(Cheng *et al*., 2008; Disterhoft *et al*., 1999; Fontan-Lozano *et al*., 2005; Weiss *et al*., 2000). To determine whether the cholinergic system modulates IN activity during TEC learning, we pharmacologically manipulated cholinergic signaling while imaging calcium transients of INs across different learning stages. The drug delivery strategy was as follows: after habituation and the first training session on day 0 (session 0), the nicotinic receptor antagonist mecamylamine (5 mg/kg, (Dai and Sun, 2024)) was administered systemically once per day, 10 minutes before training (sessions 1 to 20) from day 1 to day 4. The control group received saline injections during the same period.

As we previously reported, the conditioned response (CR %) in the mecamylamine-treated group (MEC group) significantly decreased from day 2 to day 4 compared to the saline-treated group (Saline group). Notably, blocking cholinergic signaling with MEC significantly increased SST-IN activity during the CR window but not during the unconditioned response (UR) window (Fig. 4a-c). Furthermore, the average SST-IN activity in each session showed a negative correlation with the mean EMG amplitude during the CS window (Fig. 4e). We also assessed the effect of cholinergic modulation on locomotion-related SST activity. In both running and resting trials, SST activity was higher in the MEC group than in the saline group. However, in the saline group, SST activity during running trials was lower than during resting trials, while no such difference was observed in the MEC group (Fig. 4 f,g).

**Figure 4.**
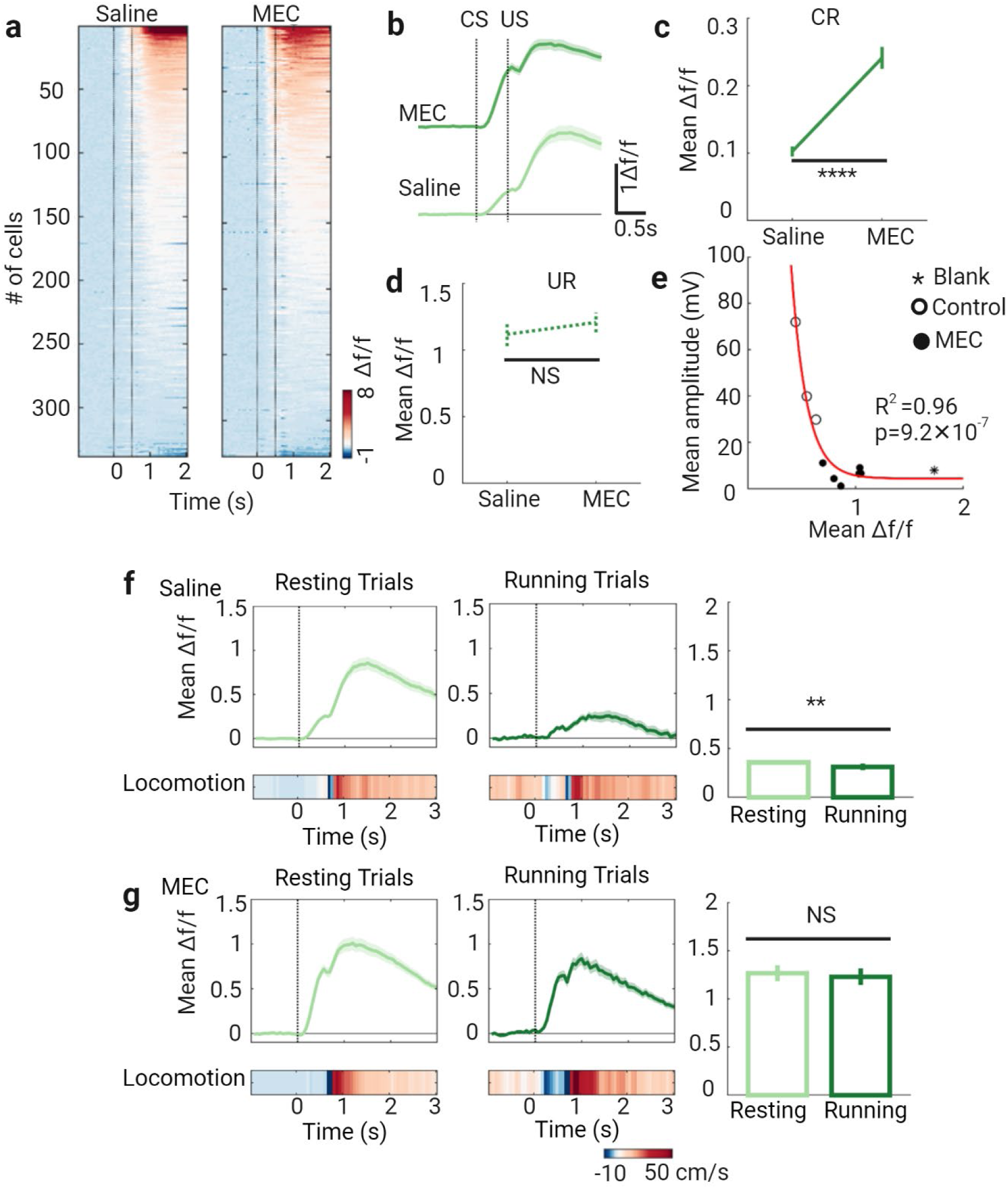
Blocking nAChRs Increases EMG- and Locomotion-Related Activity in SST Interneurons. **a.** Heat maps of GCaMP6s signals (Δf/f) for somatostatin interneurons (SST) during training sessions 1–5 in control (saline) and mecamylamine (MEC)-treated conditions. **b.** Mean signal traces of SST in saline and MEC sessions. Data are shown as mean ± s.e.m., with shaded areas indicating s.e.m. **c & d.** Response after conditioned stimulus (CS) delivery (CR, solid line, activity in CS-window) and response after unconditioned stimulus (US) delivery (UR, dashed line, activity in US-window) of SST in saline and MEC-treated sessions. **f.** Mean signal traces of SST during resting and running trials (left and middle top) in saline-treated sessions, with corresponding heatmaps of mean locomotion (left and middle bottom) and comparisons (right). Data are shown as mean ± s.e.m., with shaded areas indicating s.e.m. **g.** Same as (f) but for MEC-treated sessions.

In contrast, MEC treatment did not alter the average PV-IN activity in either the CR or UR windows, nor was there a significant relationship between PV-IN activity and EMG amplitude during the CS window (Fig. 5c). In both running and resting trials, PV activity was higher in the MEC group compared to the saline group. However, in both groups, PV activity during running trials remained lower than during resting trials (Fig 5f, g). These findings suggest that cholinergic signaling differentially modulates SST- and PV-INs, with a stronger abolishment influence on SST-INs during conditioned responses (vs. no effects on PV-INs). This highlights the distinct roles of these interneuron subtypes in processing motor-related information and shaping behavior during learning.

**Figure 5.**
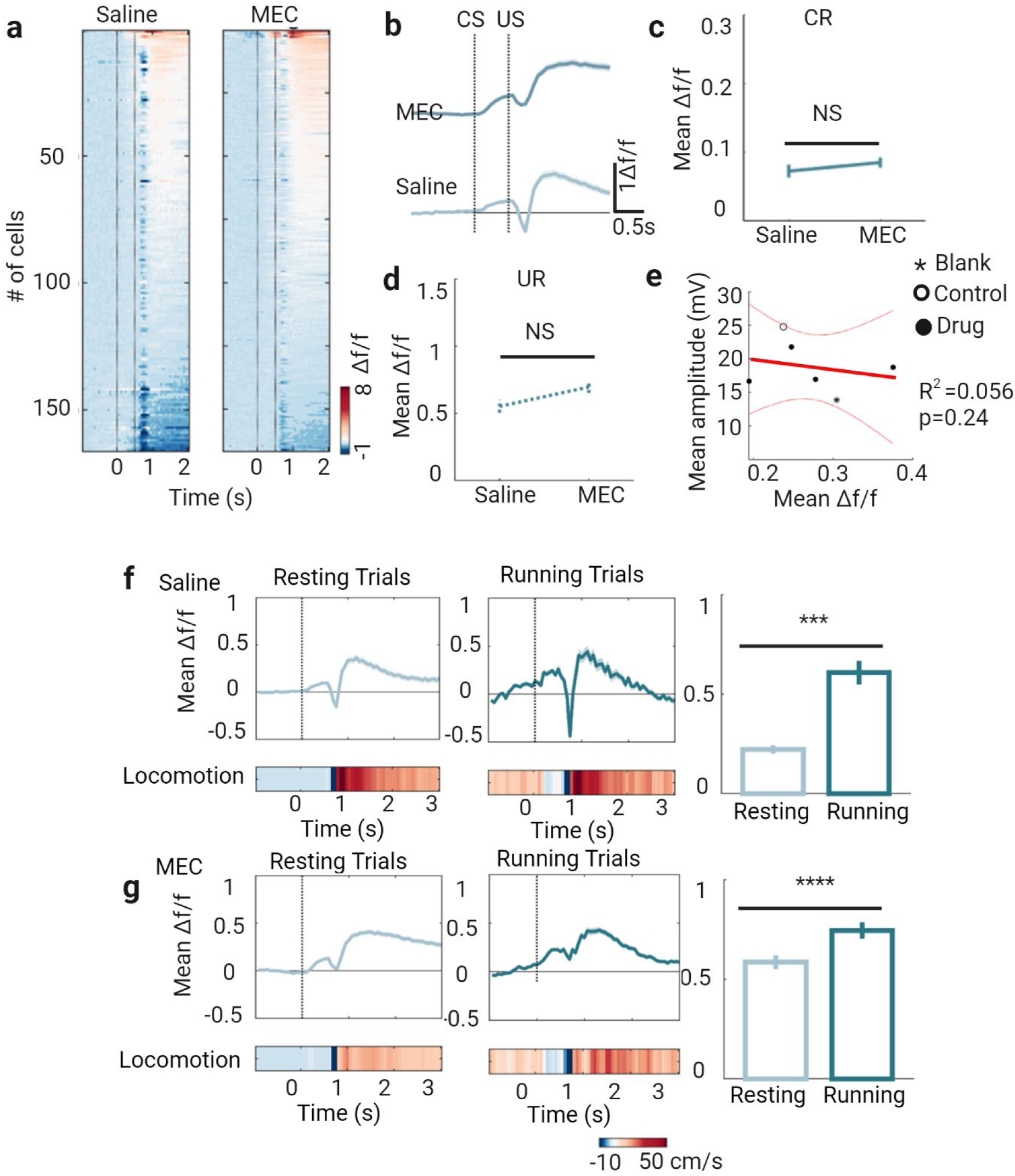
Blocking nAChRs Increases Locomotion-Modulated Activity in PV Interneurons. **a.** Heat maps of GCaMP6s signals (Δf/f) for parvalbumin interneurons (PV) during training sessions 1–5 in control (saline) and mecamylamine (MEC)-treated conditions. **b.** Mean signal traces of PV in saline and MEC sessions. Data are shown as mean ± s.e.m., with shaded areas indicating s.e.m. **c & d.** Response after conditioned stimulus (CS) delivery (CR, solid line, activity in CS-window) and response after unconditioned stimulus (US) delivery (UR, dashed line, activity in US- window) of PV in saline and MEC-treated sessions. **f.** Mean signal traces of PV during resting and running trials (left and middle top) in saline-treated sessions, with corresponding heatmaps of mean locomotion (left and middle bottom) and comparisons (right). Data are shown as mean ± s.e.m., with shaded areas indicating s.e.m. **g.** Same as (f) but for MEC-treated sessions.

## Discussion

In this study, we investigated the roles of SST-INs and PV-INs in the S1 during TEC learning. Our findings reveal distinct contributions of these interneuron subtypes to learning-related processes, particularly in modulating sensory and motor information across different stages of learning. These results highlight the complexity of inhibitory circuits in shaping behavior and learning in cortical networks and provide new insights into how interneuron diversity supports adaptive behavior.

### Dynamic Roles of SST-INs in Sensory Learning

We observed that activities of SST-INs emerge early during anticipatory eye blinks in the CS period, and that SST-INs exhibit significant changes in activity across the learning stages. Specifically, their activity increased from the habituation stage to the naive stage and decreased as animals became experts in the task. This dynamic pattern suggests that SST-INs are particularly involved in the early encoding and processing of sensory stimuli, potentially helping animals integrate and evaluate new information during the initial phases of learning. This aligns with previous studies showing that SST-INs regulate dendritic integration and plasticity, which are critical for sensory processing and learning(Karnani *et al*., 2016; Kepecs and Fishell, 2014). As learning progresses, the reduced activity of SST-INs may reflect a shift towards more refined and efficient information processing, where fewer resources are needed to process well-learned tasks. This is consistent with findings in other learning paradigms, where SST-INs modulate sensory representations during early learning stages(Khan *et al*., 2018).

### Stable yet Context-Dependent Roles of PV-INs

In contrast, PV-INs showed more stable activity during the CS window across learning stages, suggesting a consistent role in sensory processing. However, during the US window, PV-INs exhibited a similar trend to SST-INs, with activity decreasing from the naive to expert stages. This indicates that while PV-INs may not directly participate in the early phases of sensory learning, they contribute to modulating responses to the US, which might be crucial for motor control or defense mechanisms associated with the task. PV-INs are known to regulate network synchrony and gain control (Garcia-Junco-Clemente *et al*., 2019; Tremblay *et al*., 2016), and their stable activity during the CS window may reflect their role in maintaining cortical excitability during sensory processing. The differential modulation of SST- and PV-INs highlights their complementary roles in the regulation of learning and behavior, with SST-INs dynamically adapting to new information and PV-INs providing consistent inhibitory control.

### Locomotion-Dependent Modulation of Interneuron Activity

Locomotion was found to differentially affect SST- and PV-IN activity. At the habituation stage, SST-INs were less active during running trials compared to resting trials, whereas PV-INs showed increased activity during running. This suggests that SST-INs may suppress certain motor behaviors early in the learning process, while PV-INs facilitate sensory processing during movement(Garcia-Junco-Clemente *et al*., 2019). This is consistent with studies showing that locomotion enhances sensory processing in the cortex, with PV-INs playing a key role in modulating network dynamics during movement (Ayaz *et al*., 2019; Niell and Stryker, 2010). As learning progressed, the influence of locomotion on SST-IN activity diminished, while PV-INs remained more active during running trials at the naive stage but stabilized at the expert stage. These findings indicate that locomotion-dependent modulation of interneuron activity plays a dynamic role during the acquisition of new tasks, possibly reflecting a balance between sensory-motor integration and behavioral control.

### Cholinergic Modulation of Interneuron Activity

Cholinergic modulations facilitate TEC conditioning in a variety of animal models (Cheng *et al*., 2008; Disterhoft *et al*., 1999; Fontan-Lozano *et al*., 2005; Weiss *et al*., 2000) and in mice(Dai and Sun, 2024).Cholinergic signaling also played a significant role in shaping the activity of SST- and PV-INs. Pharmacological blockade of nicotinic acetylcholine receptors (nAChRs) with mecamylamine (MEC) led to increased SST-IN activity during conditioned responses (CR) but did not affect PV-IN activity. This suggests that cholinergic inputs have a more pronounced effect on SST-INs, potentially contributing to their role in processing task-relevant sensory information. This aligns with previous work showing that acetylcholine enhances cortical plasticity and sensory processing through modulation of SST-INs(Letzkus *et al*., 2011; Poorthuis *et al*., 2014). The lack of effect on PV-INs implies that their activity may be regulated by other neuromodulatory systems during learning, such as GABAergic or glutamatergic inputs (Tremblay *et al*., 2016). These results provide new insights into the role of cholinergic signaling in modulating inhibitory networks during learning and suggest that SST-INs may serve as a critical node for cholinergic regulation of sensory processing in S1 during TEC learning.

### Broader Implications and Future Directions

Our findings demonstrate that SST- and PV-INs in the primary somatosensory cortex are differentially modulated across various stages of learning and by locomotion and cholinergic signaling. SST-INs appear to play a more dynamic role in the early stages of sensory learning and are sensitive to cholinergic modulation, while PV-INs contribute more consistently to motor control and defense responses. These findings shed light on the distinct and complementary functions of interneuron subtypes in cortical circuits, advancing our understanding of how inhibitory networks support sensory processing and learning.

The differential roles of SST- and PV-INs in TEC learning have broader implications for understanding neuropsychiatric disorders characterized by interneuron dysfunction, such as schizophrenia, autism, and anxiety disorders. For example, deficits in SST-INs have been linked to impaired sensory processing in schizophrenia (Fuchs *et al*., 2017), while PV-IN dysfunction is associated with altered network synchrony in autism (Gogolla *et al*., 2009). Our findings suggest that targeting specific interneuron subtypes or cholinergic pathways could provide new therapeutic strategies for these disorders.

### Limitations of the Study

Several limitations should be acknowledged. First, the use of head-fixed mice may have influenced the behavioral responses observed, as head fixation can alter natural movement patterns and stress levels. Future studies using freely moving paradigms could provide additional insights into the role of interneurons in more naturalistic learning contexts. Second, we did not analyze the activity changes of locomotion-related and unresponsive neurons at the single-neuron level during TEC training, as these neurons are difficult to track longitudinally. Therefore, the exact origin of the UR-neurons, whether from locomotion-related or unresponsive neurons, remains unclear. Finally, the molecular identities of the CR- and UR-neurons were not explored, leaving open questions about their specific roles in learning.

## Methods

### Subjects Details

All experimental procedures were approved by the Institutional Animal Care and Use Committee (IACUC) and the Biosafety Committee at the University of Wyoming. Immune-competent male mice, aged 60 to 120 days, were used in these experiments. Mice were provided with food and water ad libitum. Before surgery, they were housed in groups of 2–5, and afterward, housed individually in a vivarium maintained at 21-23°C on a 12-hour light/dark cycle. For imaging, mice were generated by crossing Ai96 [B6;129S-Gt(ROSA)26Sortm96.1(CAG-GCaMP6s)Hze/J (JAX Stock No: 024106)] with either PV-Cre (JAX Stock No: 008069) or SST-Cre (JAX Stock No: 013044). Transgene presence (Ai96 and Cre-specific) was confirmed through PCR analysis, outsourced to Transnetyx (transnetyx.com).

### Surgeries

Mice were anesthetized with 3% isoflurane (v/v) and maintained under 2% isoflurane (v/v) in 0.4 L/min oxygen throughout the procedure. The animals were positioned on a stereotaxic frame (NARISHIGE SG-4N) and kept at 37°C using a heating pad (K&H no.1060). The incision site was sterilized with 70% isopropyl alcohol and iodine, and the scalp was shaved and cleaned. Once the skull was exposed and allowed to dry, a thin layer of Vetbond was applied over the skull and wound margins. After the adhesive dried, dental acrylic was applied around the wound edges, leaving the area over the primary somatosensory cortex (S1, right hemisphere, 1.6 mm posterior and 3.5 mm lateral to bregma) uncovered. A metal head bar was affixed caudally to the skull using dental cement, and the remaining exposed skull was sealed with additional dental cement. Mice were then placed on a heating pad for recovery. Once fully alert, they were returned to their home cage. Carprofen was administered daily for three days post-surgery, and mice were given at least seven days to recover before task training began.

To record differential EMG activity of the ipsilateral orbicularis oculi muscle, two polyimide-insulated stainless steel wires (125 μm, California Fine Wire) were implanted in the upper eyelid of the left eye, along with connecting pins. A third wire was implanted into the skull near the olfactory bulb to serve as a ground.

For two-photon imaging, a 3.5 mm circular piece of skull was removed above the primary somatosensory cortex (S1, right hemisphere, 1.6 mm posterior and 3.5 mm lateral to bregma). After removing the skull, a sterile 3 mm diameter cover glass was placed directly on the exposed dura and sealed to the surrounding skull using Vetabond.

For in vivo cannula infusion, a 1 mm craniotomy was drilled above S1 (right hemisphere, 1.6 mm posterior and 3.5 mm lateral to bregma). A guide cannula (2 mm; C315GMN/SPC; Bilaney) was then implanted and secured using Vetabond.

### Data Acquisition

#### Intrinsic Signal Imaging

To locate the C2 barrel column, intrinsic signal imaging was performed under anesthesia. The C2 whisker was inserted into a capillary connected to a piezoelectric device, which stimulated the whisker at 8 Hz (1 s pulse duration, with a 10 s gap between pulse trains), controlled by PulsePal via MATLAB. A red light source illuminated the brain surface through the cranial window, and reflectance images were captured 300 µm below the surface using a ×2.7 objective. A vasculature image was acquired using a 546-nm interference filter and superimposed onto the intrinsic signal image.

#### Two-Photon Calcium Imaging

Two-photon calcium imaging was performed using a resonant/galvo scanning two-photon microscope (Neurolabware, Los Angeles, CA) controlled by Scanbox software(Yaeger *et al*., 2019). GCaMP6s was excited at 920 nm using an Insight X3 laser (Spectra-Physics, Milpitas, USA), focused through a 16×/0.8NA water-immersion objective. Images (512 x 796 pixels, covering a 490 x 630 μm field) were acquired at a 15.5 Hz frame rate at depths of 120 to 320 μm below the pial surface for layer 2/3 imaging. During imaging, mice moved freely on a 3D-printed running wheel, and locomotion was recorded using a rotary encoder, synchronized with the imaging frame rate. TTL signals for whisker stimulation and air puff, controlled by PulsePal, were used to align the two-photon imaging data.

#### EMG Acquisition

Electromyography (EMG) signals from the eyelid were band-pass filtered (50–500 Hz), amplified using a 4-channel differential amplifier (model 1700, A-M Systems), and acquired via a 16-bit data acquisition system (Digidata 1322A, Axon Instruments), sampled at 1000 Hz. TTL signals for whisker stimulation and air puff, output from PulsePal, were used to synchronize EMG signals with the stimulus delivery.

#### Stimulation Delivery

Whisker stimulation was delivered using an E-650 Piezo Amplifier, controlled by PulsePal. An air puff to the eye was administered via a solenoid valve (PSV-5, Aalborg) controlled by PulsePal. The entire two-photon imaging system, along with whisker stimulation, air puff, and laser stimuli, was controlled by custom software written in MATLAB.

### Behavioral Procedures

Behaviors were performed according to a procedure described earlier(Dai and Sun, 2024). Briefly, prior to training, mice were habituated to head restraint for 3 days. During this period, they were placed on the running wheel with their heads fixed for 1 hour each day. From day 4 to day 6, whisker stimulation (10 Hz) was delivered in 4 sessions (30 trials per session) while two-photon imaging was performed based on the results of intrinsic signal imaging. On day 7, the whisker stimulation frequency was increased to 60 Hz in a single session (30 trials), referred to as the "Habituation stage" (Hab) in this study.

Following habituation, trace eyeblink conditioning (TEC) was initiated. The conditioned stimulus (CS) was whisker deflection (60 Hz, 250 ms), followed by an unconditioned stimulus (US) after a 250 ms interval (from CS offset to US onset). The US was a periorbital air puff (15 psi, 50 ms), delivered through a plastic needle positioned 5 mm from the cornea. The interval between trials varied randomly from 20 to 40 seconds. Each mouse underwent 5 consecutive sessions per day (20 trials per session) until it learned to maintain eyelid closure from CS onset until US offset, as determined by EMG analysis (details in Data Analysis). The first training session was referred to as the "Naive stage" (Naive), and the final session was called the "Expert stage" (Exp). In each session, 80% of trials involved paired CS and US (CS-US), 10% were CS-only trials, and 10% were US-only trials, with trial order randomized. If a mouse showed signs of discomfort, it was returned to its home cage and given up to 24 hours to recover.

### Pharmacological manipulation of cholinergic inputs

Pharmacological manipulations were performed according to a procedure described earlier(Dai and Sun, 2024). All drug or saline injections began from the second training session. Injections were administered at the start of each day, with a 10-minute waiting period before resuming the TEC training (5 sessions per day). Drugs were dissolved in physiological saline (0.9% sodium chloride). For local inactivation, mecamylamine (0.5 µg/µl) was delivered through the implanted cannula via pressure injection, using a custom device controlled by a single-axis hydraulic manipulator (NARISHIGE MMO-220A) at a rate of 30–50 nL/min.

### Data processing and analysis

The two-photon image processing pipeline followed previously established methods. Briefly, movies from the same plane for each session were concatenated and motion-corrected. After alignment, the mean image was generated to track the same cells for longitudinal imaging. Segmentation was performed using a MATLAB graphical user interface tool (Scanbox, Los Angeles, CA) to define regions of interest (ROIs) corresponding to neuronal somas. Calcium transients for each cell were extracted, and estimated spiking was calculated using non-negative temporal deconvolution of the corrected ROI signal with the Vanilla algorithm.

For calcium transient analysis, we calculated the baseline signal (∼930 frames, 60 seconds before conditional stimulus (CS) onset) for each cell by selecting the minimum of the mean values from two segments (1–450 frames and 451–900 frames). The ratio of Δf/f at each time point was then computed. Each session’s signal was segmented into 30 trials (Habituation stage) or 20 trials (Naive and Expert stages), aligned by the onset of the CS (1 second before to 2 seconds after CS onset).

For locomotion analysis, we similarly calculated the baseline locomotion signal (∼930 data points, 60 seconds before CS onset) by selecting the minimum mean values from the same two segments. The baseline was subtracted, and the locomotion traces were segmented into trials aligned by the CS onset.

For electromyography (EMG) analysis, the EMG data was resampled to match the length of the locomotion or calcium transient data based on the TTLs of CS and unconditioned stimulus (US). The baseline EMG signal (∼930 data points, 60 seconds before CS onset) was established and subtracted in the same manner. EMG traces were segmented into trials aligned by the CS onset. The conditioned response (CR) was defined as EMG activity exceeding 4 standard deviations above the baseline for at least 200 milliseconds before US onset. A CR percentage (CR %) greater than 60% within a session indicated that the learning criterion was met. The early CS response (ECR) represented the mean amplitude of EMG from CS onset to CS offset, while the late CS response (LCR) was the average amplitude from CS offset to US onset. The US response (UR) was the mean amplitude of EMG recorded for 500 milliseconds after US onset.

Linear regression analysis was performed using MATLAB’s ‘fitlm’ function. For decay function fitting, the following equation was used:

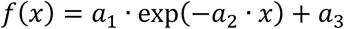

Here, *x* is the input, and *a*_1_, *a*_2_, *a*_3_ are the fitting parameters. The MATLAB function ‘fitnlm’ was then employed to fit the data.

### Statistical analysis

Statistical details are provided where applicable. Data are expressed as mean ± standard error of the mean (s.e.m.). Analyses were performed using MATLAB. Parametric tests were applied whenever possible to compare means. Non-parametric tests were employed for non-normally distributed data. Two-tailed tests were used unless otherwise specified. A p-value of less than 0.05 was considered statistically significant.

## Acknowledgements

We thank Dr. Z. Zhang for imaging and C. Zhang for animal husbandry, histology assistance and items purchasing. We thank Dr. Y. Li and Y. Wang for discussion. This work is supported by grants from National Institute of Mental Health (1R21MH131363), National Institute of Biomedical Imaging and Bioengineering (1R21EB032609), National Institute on Aging (5R21AG072803) and from National Institute of General Medical Sciences (2P20GM121310) and administrative supplements from National Institute of Aging (3P20GM121310-07S3) and Office of The Director, National Institutes Of Health (OD, 3P20GM121310-07S2).

## Author contributions

Q.Q.S. designed the experiment and supervised the project, acquired the funding, and wrote the manuscript. J.D. built the behavioral set-up, co-designed the experiments, collected and analyzed data, and wrote the manuscript.

## References

Ayaz, A., Stauble, A., Hamada, M., Wulf, M. A., Saleem, A. B. and Helmchen, F. (2019). Layer-specific integration of locomotion and sensory information in mouse barrel cortex. Nat Commun, 10, 2585.

Cheng, D. T., Disterhoft, J. F., Power, J. M., Ellis, D. A. and Desmond, J. E. (2008). Neural substrates underlying human delay and trace eyeblink conditioning. Proc Natl Acad Sci U S A, 105, 8108–8113.

Christian, K. M. and Thompson, R. F. (2005). Long-term storage of an associative memory trace in the cerebellum. Behav Neurosci, 119, 526–537.

Dai, J. and Sun, Q. Q. (2024). Modulation of cortical representations of sensory and contextual information underlies aversive associative learning. Cell Rep, 43, 114672.

Del Rosario, J., Coletta, S., Kim, S. H., Mobille, Z., Peelman, K., Williams, B., Otsuki, A. J., Del Castillo Valerio, A., Worden, K., Blanpain, L. T., Lovell, L., Choi, H. and Haider, B. (2025). Lateral inhibition in V1 controls neural and perceptual contrast sensitivity. Nat Neurosci.

Disterhoft, J. F., Kronforst-Collins, M., Oh, M. M., Power, J. M., Preston, A. R. and Weiss, C. (1999). Cholinergic facilitation of trace eyeblink conditioning in aging rabbits. Life Sci, 64, 541–548.

Felleman, D. J. and Van Essen, D. C. (1991). Distributed hierarchical processing in the primate cerebral cortex. Cereb Cortex, 1, 1–47.

Ferguson, K. A., Salameh, J., Alba, C., Selwyn, H., Barnes, C., Lohani, S. and Cardin, J. A. (2023). VIP interneurons regulate cortical size tuning and visual perception. Cell Rep, 42, 113088.

Fontan-Lozano, A., Troncoso, J., Munera, A., Carrion, A. M. and Delgado-Garcia, J. M. (2005). Cholinergic septo-hippocampal innervation is required for trace eyeblink classical conditioning. Learn Mem, 12, 557–563.

Freedman, D. J. and Assad, J. A. (2016). Neuronal Mechanisms of Visual Categorization: An Abstract View on Decision Making. Annu Rev Neurosci, 39, 129–147.

Fuchs, T., Jefferson, S. J., Hooper, A., Yee, P. H., Maguire, J. and Luscher, B. (2017). Disinhibition of somatostatin-positive GABAergic interneurons results in an anxiolytic and antidepressant-like brain state. Mol Psychiatry, 22, 920–930.

Garcia-Junco-Clemente, P., Tring, E., Ringach, D. L. and Trachtenberg, J. T. (2019). State-Dependent Subnetworks of Parvalbumin-Expressing Interneurons in Neocortex. Cell Rep, 26, 2282–2288 e2283.

Giovannucci, A., Badura, A., Deverett, B., Najafi, F., Pereira, T. D., Gao, Z., Ozden, I., Kloth, A. D., Pnevmatikakis, E., Paninski, L., De Zeeuw, C. I., Medina, J. F. and Wang, S. S. (2017). Cerebellar granule cells acquire a widespread predictive feedback signal during motor learning. Nat Neurosci, 20, 727–734.

Gogolla, N., Leblanc, J. J., Quast, K. B., Sudhof, T. C., Fagiolini, M. and Hensch, T. K. (2009). Common circuit defect of excitatory-inhibitory balance in mouse models of autism. J Neurodev Disord, 1, 172–181.

Heiney, S. A., Wohl, M. P., Chettih, S. N., Ruffolo, L. I. and Medina, J. F. (2014). Cerebellar-dependent expression of motor learning during eyeblink conditioning in head-fixed mice. J Neurosci, 34, 14845–14853.

Karnani, M. M., Jackson, J., Ayzenshtat, I., Hamzehei Sichani, A., Manoocheri, K., Kim, S. and Yuste, R. (2016). Opening Holes in the Blanket of Inhibition: Localized Lateral Disinhibition by VIP Interneurons. J Neurosci, 36, 3471–3480.

Kepecs, A. and Fishell, G. (2014). Interneuron cell types are fit to function. Nature, 505, 318–326.

Khan, A. G., Poort, J., Chadwick, A., Blot, A., Sahani, M., Mrsic-Flogel, T. D. and Hofer, S. B. (2018). Distinct learning-induced changes in stimulus selectivity and interactions of GABAergic interneuron classes in visual cortex. Nat Neurosci, 21, 851–859.

Kvitsiani, D., Ranade, S., Hangya, B., Taniguchi, H., Huang, J. Z. and Kepecs, A. (2013). Distinct behavioural and network correlates of two interneuron types in prefrontal cortex. Nature, 498, 363–366.

Letzkus, J. J., Wolff, S. B., Meyer, E. M., Tovote, P., Courtin, J., Herry, C. and Luthi, A. (2011). A disinhibitory microcircuit for associative fear learning in the auditory cortex. Nature, 480, 331–335.

Makino, H., Hwang, E. J., Hedrick, N. G. and Komiyama, T. (2016). Circuit Mechanisms of Sensorimotor Learning. Neuron, 92, 705–721.

Marín, O. (2012). Interneuron dysfunction in psychiatric disorders. Nature Reviews Neuroscience, 13, 107–120.

McEchron, M. D., Bouwmeester, H., Tseng, W., Weiss, C. and Disterhoft, J. F. (1998). Hippocampectomy disrupts auditory trace fear conditioning and contextual fear conditioning in the rat. Hippocampus, 8, 638–646.

Moyer, J. R., Jr., Deyo, R. A. and Disterhoft, J. F. (1990). Hippocampectomy disrupts trace eye-blink conditioning in rabbits. Behav Neurosci, 104, 243–252.

Myers-Joseph, D., Wilmes, K. A., Fernandez-Otero, M., Clopath, C. and Khan, A. G. (2024). Disinhibition by VIP interneurons is orthogonal to cross-modal attentional modulation in primary visual cortex. Neuron, 112, 628–645 e627.

Niell, C. M. and Stryker, M. P. (2010). Modulation of visual responses by behavioral state in mouse visual cortex. Neuron, 65, 472–479.

Pfeffer, C. K., Xue, M., He, M., Huang, Z. J. and Scanziani, M. (2013). Inhibition of inhibition in visual cortex: the logic of connections between molecularly distinct interneurons. Nat Neurosci, 16, 1068–1076.

Poort, J., Khan, A. G., Pachitariu, M., Nemri, A., Orsolic, I., Krupic, J., Bauza, M., Sahani, M., Keller, G. B., Mrsic-Flogel, T. D. and Hofer, S. B. (2015). Learning Enhances Sensory and Multiple Non-sensory Representations in Primary Visual Cortex. Neuron, 86, 1478–1490.

Poorthuis, R. B., Enke, L. and Letzkus, J. J. (2014). Cholinergic circuit modulation through differential recruitment of neocortical interneuron types during behaviour. J Physiol, 592, 4155–4164.

Szadai, Z., Pi, H. J., Chevy, Q., Ocsai, K., Albeanu, D. F., Chiovini, B., Szalay, G., Katona, G., Kepecs, A. and Rozsa, B. (2022). Cortex-wide response mode of VIP-expressing inhibitory neurons by reward and punishment. Elife, 11.

Tremblay, R., Lee, S. and Rudy, B. (2016). GABAergic Interneurons in the Neocortex: From Cellular Properties to Circuits. Neuron, 91, 260–292.

Weiss, C., Preston, A. R., Oh, M. M., Schwarz, R. D., Welty, D. and Disterhoft, J. F. (2000). The M1 muscarinic agonist CI-1017 facilitates trace eyeblink conditioning in aging rabbits and increases the excitability of CA1 pyramidal neurons. J Neurosci, 20, 783–790.

Woodruff-Pak, D. S. (1993). Eyeblink classical conditioning in H.M.: delay and trace paradigms. Behav Neurosci, 107, 911–925.

Yaeger, C. E., Ringach, D. L. and Trachtenberg, J. T. (2019). Neuromodulatory control of localized dendritic spiking in critical period cortex. Nature, 567, 100–104.

